# Feedforward and feedback frequency-dependent interactions in a large-scale laminar network of the primate cortex

**DOI:** 10.1101/065854

**Authors:** Jorge F. Mejias, John D. Murray, Henry Kennedy, Xiao-Jing Wang

## Abstract

Interactions between top-down and bottom-up processes in the cerebral cortex hold the key to understanding predictive coding, executive control and a gamut of other brain functions. The underlying circuit mechanism, however, remains poorly understood and represents a major challenge in neuroscience. In the present work we tackled this problem using a large-scale computational model of the primate cortex constrained by new directed and weighted connectivity data. In our model, the interplay between feedforward and feedback signaling depends on the cortical laminar structure and involves complex dynamics across multiple (intra-laminar, inter-laminar, inter-areal and whole cortex) scales. The model was tested by reproducing, and shedding insights into, a wide range of neurophysiological findings about frequency-dependent interactions between visual cortical areas: feedforward pathways are associated with enhanced gamma (30-70 Hz) oscillations, whereas feedback projections selectively modulate alpha/low beta (8-15 Hz) oscillations. We found that in order for the model to account for the experimental observations, the feedback projection needs to predominantly target infragranular layers in a target area, which leads to a proposed circuit substrate for predictive coding. The model reproduces a functional hierarchy based on frequency-dependent Granger causality analysis of inter-areal signaling, as reported in recent monkey and human experiments. Taken together, this work highlights the importance of multi-scale approaches and provides a modeling platform for studies of large-scale brain circuit dynamics and functions.

## 1 Introduction

Inasmuch as the primate cerebral cortex is organized hierarchically, it is essential to understand interactions between feedforward (bottom-up) information processing and feedback (top-down) signaling which may mediate brains prediction about the sensory world, attention, behavioral context and control. With the advance of electrode recording arrays and other techniques, experimental studies in recent years not only have yielded increasing information about dynamical interactions between pairs of cortical areas, but also begun to reveal the complex feedforward and feedback signaling flow among many cortical regions in a large-scale system. Motivated by new experimental observations, in this work we built an anatomically-based large-scale model of the primate cortex endowed with a laminar structure, and used it to elucidate the dynamical interplay between feedforward and feedback signals at the global brain level.

In the macaque visual cortical system, an anatomical hierarchy is characterized by specific laminar patterns of feedforward and feedback inter-areal projections [1, 2]. Feedforward (FF) projections tend to originate from supragranular layers and target layer 4 [1, 2], whereas feedback (FB) projections largely stem from infragranular layers and target supra- and infragranular layers while avoiding layer 4 [1, 2]. In addition, the laminar restriction of these projections increases with the hierarchical distance: if two areas are widely separated, the supragranular origin of FF projections are more pronounced than if the areas are closer to each other in the hierarchy, and the same holds true for the infragranular origin of FB projections [3, 2].

Hierarchical interactions between visual cortical areas occur in both FF and FB directions, with FF interactions transmitting sensory information from lower to higher brain areas and FB interactions conveying top-down modulation to early sensory areas [4]. Recently, it has been observed that these interactions are characterized by distinct neural oscillatory patterns: interactions in the FF direction are associated with enhanced gamma oscillations (30-70 Hz), while FB interactions are associated with enhanced alpha or low beta oscillations (8-15 Hz) [5, 4, 6, 7]. This frequency profile has been observed in macaque using different recording techniques (from multi-contact electrodes [4] to ECoG grids [6]) and, more recently, in human using magnetoencephalography [7]. It is plausible that these enhanced gamma/alpha signatures in FF/FB interactions may be related to the laminar preference of the anatomical projections described above [8]. However, it is unclear what the neural circuit principles are through which oscillatory dynamics originating within local microcircuits drive the spatially segregated, frequency-specific interactions within dense large-scale networks.

To address this question, we built a large-scale computational model of laminar cortical microcircuits with 30 cortical areas that interact via their long-range cortico-cortical feedforward and feedback pathways. This large-scale cortical network was built on anatomical inter-areal connectivity data from tract tracing studies [9, 10, 2]. The connectivity data that we present here extends previous studies [3, 2] by including novel data on the parietal subnetwork (area LIP), yielding a large-scale network of 30 cortical areas distributed across the occipital, temporal, parietal and frontal lobes. This rich quantitative dataset contains information on the directed strength of inter-areal projections, their laminar origin, and the propagation latency obtained from the wiring distances between areas [11]. We incorporated dynamics into the large-scale cortical circuit by modeling each local circuit with interacting excitatory and inhibitory populations, so that supragranular and infragranular layers differentially exhibit fast (gamma) and slow (alpha-beta) neural oscillatory dynamics, respectively.

The model spans four spatial scales: intra-laminar, inter-laminar, inter-areal, and large-scale levels (see Fig. 1 for a schematic representation). At each level, we constrained the circuit using empirical anatomical data, comparing the computed dynamics with published electrophysiological findings and providing novel and valuable insight about cortical circuit mechanisms. The parsimonious model is validated by capturing a wide range of electrophysiological observations, including (i) the predominance of weakly coherent gamma rhythms in supra-and strongly coherent alpha in infragranular layers [12], (ii) the effects of visual contrast on gamma frequency and power in V1 [13, 14, 15], (iii) the inter-laminar entrainment between supragranular gamma power and infragranular alpha phase [16], (iv) the inverse correlation between alpha power and supragranular firing rates, which associates strong alpha rhythms with local inhibition [17, 18], and (v) the evidence of enhanced gamma and alpha signatures in FF and FB inter-areal interactions [5, 4, 6]. Altogether, the model provides insight into plausible mechanisms underlying top-down attention signals and predictive coding [19].

**Figure 1:**
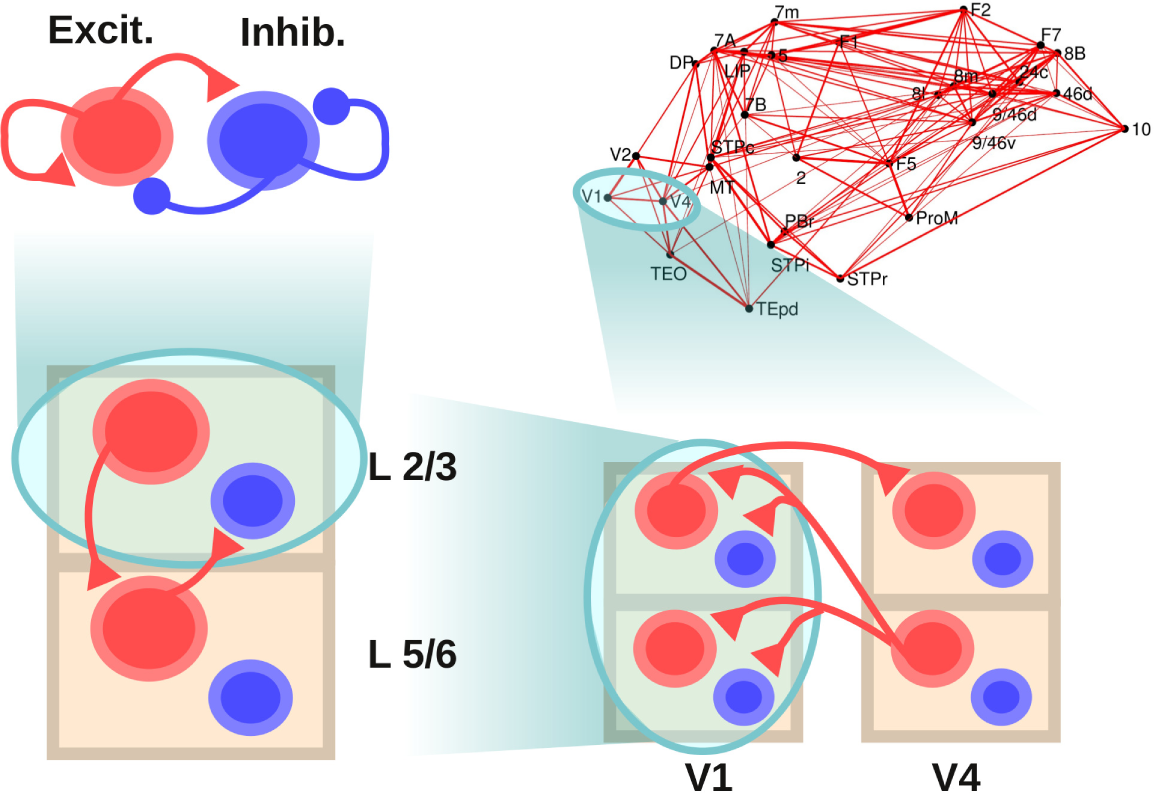
Scheme of the large-scale model. The scheme shows the four levels considered: a within-layer, local microcircuit consisting of an excitatory (in red) and an inhibitory (in blue) population (upper left), a laminar circuit with two laminar modules (corresponding to supragranular and infragranular layers, lower left), an interareal circuit with laminar-specific projections (lower right), and a large-scale network of 30 cortical areas based on macaque anatomical connectivity (upper right). Each level is anatomically constrained, and its dynamics provides insight at different electrophysiological observations in macaque. Only the connections at each level not shown at a lower level are plotted, for clarity.

With laminar circuit models embedded within each area, and interacting through the structured inter-areal connectivity, our main finding is that the large-scale model captures the laminar-specific feedforward and feedback interactions on a global scale across many areas. It quantitatively predicts the emergence of frequency-dependent functional interactions and its relationship to underlying anatomical connectivity [6, 7]. Importantly, it provides a mechanistic explanation for the emergence of a functional hierarchy among visual cortical areas, as recently observed in macaques [6] and humans [7], and it allows exploration of flexible nature of brains hierarchy which could be altered by behavioral context. Our work highlights the importance of multi-scale approaches in the construction of large-scale brain models.

## 2 Results

The model spans four spatial levels of description, as depicted in Fig. 1: intra-laminar, inter-laminar, inter-areal, and large-scale network. Each level is constrained with anatomical and electrophysiological data, and forms the basis upon which the next level is built. We report results of this work across the four levels sequentially.

### 2.1 Intra-laminar level

We first consider the local or intra-laminar microcircuit, which is the lowest level of the model and may be identified as a set of neurons within a given cortical area and layer. More precisely, we assume at this level a population of pyramidal neurons and a population of inhibitory interneurons. Within and between each population are recurrent and cross connections, respectively (Fig. 2A, top). Local strongly interconnected populations of excitatory and inhibitory neurons are present in both supra- and infragranular cortical layers [20], and their dynamics have been extensively studied using diverse computational approaches [21, 22, 23]. For this level of description, we simulate each laminar sub-circuit with a nonlinear firing-rate model, of the Wilson-Cowan type (see Methods), which represents the mean activities of a population of excitatory neurons and a population of inhibitory neurons.

**Figure 2:**
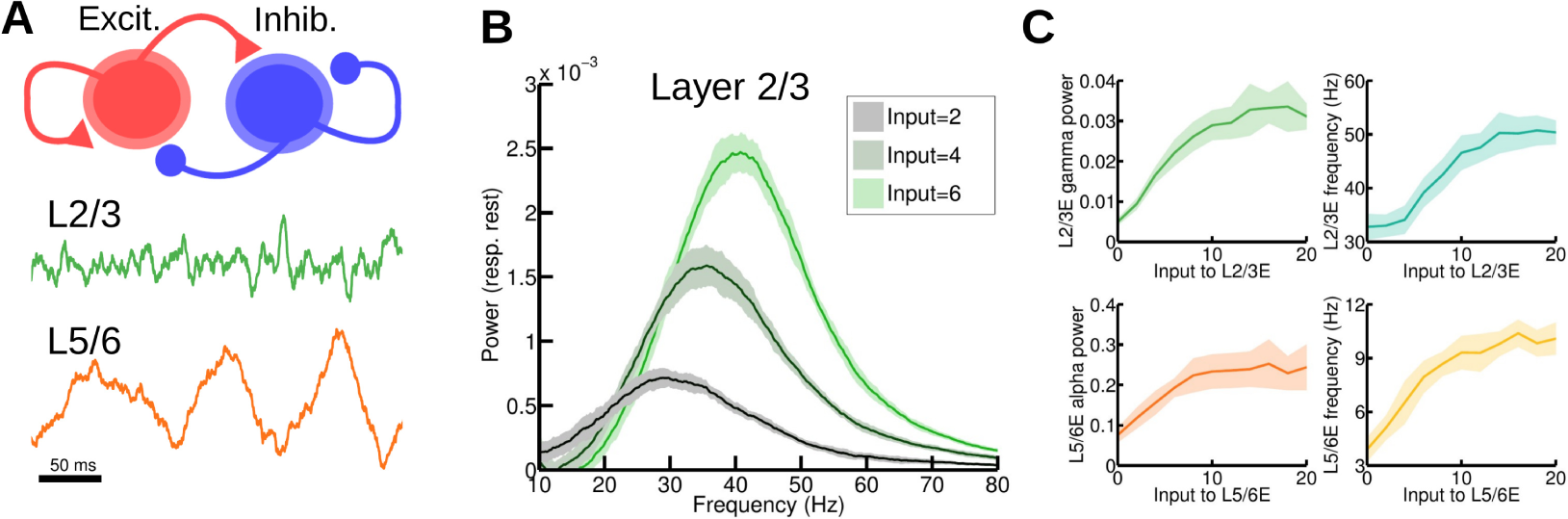
Local circuit model at the intra-laminar level. (A) Scheme of the local circuit (top), with the excitatory and inhibitory population in red and blue, respectively, and examples of the oscillatory activity for an excitatory-inhibitory (E-I) circuit in layer 2/3 (middle, in green) and layer 5/6 (bottom, in orange). (B) Power spectrum of the firing rate of an isolated layer 2/3 as a function of input strength to the excitatory population. The spectrum of the spontaneous state (with zero input) has been subtracted in each case to highlight changes induced by the input (see main text). As the input increases (which resembles the effect of increasing the contrast of a visual stimulus) the power of gamma rhythms becomes stronger, as in observations by Henrie and Shapley. (C) Effect of the input to the excitatory population on the power spectrum peak (left) and frequency (right) of the oscillations, for an isolated layer 2/3 (top) and an isolated layer 5/6 (bottom).

The local circuits of the supra- and infragranular layers differ in their connectivity and physiology, leading to their distinct dynamics [24, 20]. For the supragranular circuits, we constrained parameter values so that the populations display a noise-driven gamma (~40 Hz) rhythm (Fig. 2A, middle), as is commonly observed in layer 2/3 [25, 12, 16, 4]. For the infragranular circuits, we adjusted parameter values so that the oscillations displayed by the model fall within the alpha (~10 Hz) or low beta (~15-30 Hz) frequency range [12, 4] (Fig. 2A, bottom).

A simple coupled excitatory-inhibitory system as described here is useful for studying the response of early visual neurons to incoming visual stimuli. For instance, local field potential (LFP) shows that increasing the contrast of a visual grating enhances the strength of gamma activity in macaque V1 for low and medium contrast levels [13, 15]. A higher visual contrast has also been associated with higher frequencies of the gamma activity [14], while other factors such as the stimulus size, the level of masking noise and the stimulus orientation likewise have an effect on gamma power and frequency [15].

In order to test the behavior of the model at the intra-laminar level, we simulated the effect of an increase of stimulus contrast on gamma rhythms for layer 2/3 neurons in V1, and compared the outcome on those observed experimentally [13]. We modeled the increase of visual contrast as an increase of the input to the excitatory population, since increases in contrast of a stimulus falling within the receptive field of a neuron elicit increases in firing rate [13]. As shown in Fig. 2B, higher contrast values lead to stronger gamma rhythms, characterized by a higher gamma peak of the excitatory firing rate power spectrum. Note that, as in [13], the power spectrum corresponding to spontaneous activity (i.e. zero input) was subtracted from the curves showed, thereby removing the power-law appearance of the power spectra but nevertheless conserving the effects of the input in a principled fashion. The enhancement of the power spectrum is strongest on the 30~60 Hz gamma band, as in [13] (cf. its figure 4). The contrast-mediated enhancement of the high gamma (>100 Hz) rhythm observed experimentally is not fully captured by our model. This is to be expected given that the broadband power at this range reflects individual spiking activity [26, 27, 28], and a rate model will fail to reproduce this feature. This does not constitute a problem, however, since we are not considering effects on high gamma rhythms (i.e. larger than 80 Hz) in this study.

A second effect shown in Fig. 2B is a small but consistent shift of the gamma peak towards higher frequencies (from 30 to 40 Hz, approximately) with increasing contrast. Although not observed in the study from Henrie and Shapley [13], this effect of contrast on gamma frequency has been consistently found in more recent studies [28, 15], and the model captures this effect. Fig. 2C shows in more detail the influences of increasing the input to the excitatory population on the peak power (left) and frequency (right) of the oscillations, both for the case of an isolated layer 2/3 microcircuit (gamma rhythm, top) and an isolated layer 5 microcircuit (alpha rhythm, bottom). Note that the power and frequency curves saturate at high inputs, in line with evidence of nonlinear effects on V1 gamma power for strong contrast [15, 29].

The local circuit presented here displays, therefore, the same increases of power and frequency of neural oscillations with increasing input that are observed in recent electrophysiological studies of early visual areas. This makes the circuit a good starting point to understand rhythmic interactions in larger neural systems.

### 2.2 Inter-laminar level

Having characterized the neural dynamics of an isolated layer, we built a laminar circuit by considering several layers and adding inter-laminar projections between them. In order to investigate the interplay between gamma and alpha/low beta rhythms, we consider two laminar modules, each one of them as a generator of the gamma and alpha rhythm, respectively. Gamma oscillations are most prominently found in granular and supragranular layers before propagating to other layers [12, 16, 4], and modeling work suggest that layers 2/3 and 4 may locally generate gamma rhythms [23]. On the other hand, slower oscillations in the alpha and low beta frequency range are most strongly present at infragranular layers, from where they propagate to other layers [4]. We therefore assume two laminar modules: a supragranular module (layer 2/3) that displays gamma rhythms, and an infragranular module (layers 5 and 6) that displays alpha rhythms.

In order to couple the modules of both layers, for parsimony we consider a subset of strong anatomical projections in each direction as a first approximation of the inter-laminar circuit. Our hypothesis is that, in the context of gamma-alpha rhythm interactions, the dynamics of the circuit may be described reasonably well by considering uniquely these strong projections, while further details add more richness and complexity to the inter-laminar dynamics.

Anatomical studies indicate that, in the supragranular-to-infragranular direction, there is a predominant excitatory projection from layer 3 pyramidal neurons to layer 5 pyramidal neurons, which is consistently stronger than most other inter-laminar projections [20, 24]. In the infragranular-to-supragranular direction, there is a particularly strong projection from layer 5 pyramidal neurons to layer 2/3 interneurons [30, 31]. Comparative studies have confirmed these two projections as the strongest ones between layers 2/3 and 5, with strong projections from layer 6 pyramidal neurons to layer 4 inhibitory cells (which later project to layer 2/3 pyramidal cells) further supporting the inhibitory role of infragranular-to-supragranular projections [32, 33]. We therefore set these two projections (L2/3E-to-L5E and L5E-to-L2/3I) as our main inter-laminar connections in the model, as shown in Fig. 3A (left).

**Figure 3:**
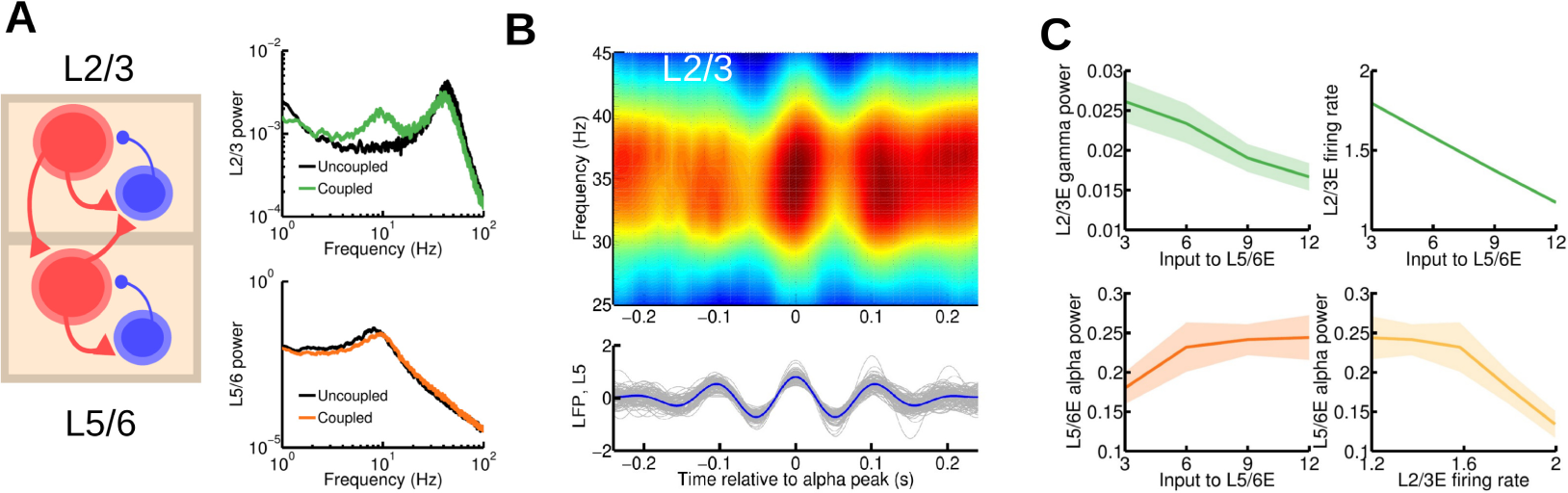
Cortical area model at the inter-laminar level. (A) Scheme of the inter-laminar circuit (left panel), self-connections within a given population are omitted in the figure for clarity. Inter-laminar connections considered in the model correspond to the strongest projections between layer 2/3 and layer 5/6 as found in experimental studies. Right panel: power spectrum of layer 2/3 (top) and 5/6 (bottom) in the case of uncoupled, isolated layers (in black, for comparison) and interconnected network (green and orange, respectively). A background input of I=8 was fed into the excitatory population of both layers. (B) Lower panel: a set of 30 traces of activity in layer 5/6 (in gray) and their average (in blue). The central peak of each trace was aligned at zero before averaging. Upper panel: a periodogram of layer 2/3 showing the averaged power for a range of frequencies for the same temporal periods as the layer 5/6 traces. We can see the existence of a strong entrainment of gamma power to alpha phase, as in the experimental findings by Spaak *et al.* Input was I=6 for supragranular and I=8 for infragranular excitatory populations. (C) Effect of injecting external current to the excitatory population of layer 5/6 on the layer 2/3 gamma power and (dimensionless) firing rate (top left and right, respectively) and on layer 5/6 alpha power (bottom left). An inverse relationship between supragranular firing rate and alpha power is observed (bottom right), which highlights a possible link of enhanced alpha rhythms with activity suppression.

As a result of the inter-laminar coupling, rhythms spread across layers and the oscillatory dynamics in both supra- and infragranular layers presents a more dynamically rich profile, compared to the isolated layers. As Fig. 3A (right) shows, the power spectrum of layer 2/3 activity now displays a strong alpha component as a consequence of the oscillatory input coming from layer 5/6. The effect of layer 2/3 activity on the dynamics of layer 5/6 is slightly different, however: because of the intrinsically slower dynamics of layer 5/6, the fast fluctuations due to the weak gamma rhythm are partially filtered and the input from layer 2/3 to layer 5/6 is a quasi-constant term that shifts the alpha peak towards higher frequencies.

The alpha-modulated input arriving at layer 2/3 from layer 5/6 also has a powerful impact, leading to the entrainment of gamma and alpha rhythms (Fig. 3B). Here, the slow oscillatory input from layer 5/6 modulates the power of gamma oscillations in layer 2/3, leading to a phase-amplitude coupling (PAC) between layer 2/3 gamma and layer 5/6 alpha rhythms. This PAC phenomenon has been observed in multielectrode recordings in macaque ([16], cf. its figure 2), and constitutes a validation of our model at the inter-laminar level.

To further characterize the behavior of the multi-laminar circuit, we studied the effects of external input on its dynamics. The case of an external constant input to layer 5/6 is particularly interesting (Fig. 3C). A constant input arriving at the excitatory population of layer 5/6 has two main effects on layer 2/3, due to inputs to the inhibitory neurons in layer 2/3: a decrease in gamma power and a decrease in mean firing rate of layer 2/3 pyramidal cells (top panels in Fig. 3C). In addition, the input to layer 5/6 enhances the infragranular alpha rhythm (bottom left panel), as observed for the isolated layer case in Fig. 2. This reveals a significant negative correlation between supragranular mean firing rate and alpha power (bottom right panel), which strongly supports the idea that alpha rhythms reflect local inhibition of areas not involved in a particular cognitive task [18].

In summary, our laminar model of cortical area consisting on two laminar modules (for supragranular and infragranular) constrained by anatomical data displays phase-amplitude coupling between gamma and alpha rhythms as observed experimentally, and provides insight on a possible relationship between alpha rhythms and activity suppression.

### 2.3 Inter-areal level

Extending our description to the inter-areal level involves characterizing the anatomical projections linking the microcircuits of two or more cortical areas. Here, we assume two areas with a clearly defined hierarchical relationship (for example, areas V1 and V4). FF projections along the visual hierarchy stem from supragranular layers and preferentially target layer 4, which in turn projects to layer 2/3 within that area [1, 2]. In our model, this is approximated as a projection from layer 2/3 pyramidal neurons in V1 to layer 2/3 pyramidal neurons in V4. By contrast, FB projections originate from infragranular layers (predominantly from layer 6) and target supra- and infragranular layers while avoiding layer 4 [1, 2]. Recent findings indicate that FB projections from these layers are more diffuse than FF in terms of their targets [2]. In our model, we assume that the FB projection stems from layer 5/6 pyramidal neurons in V4 and targets all four populations in V1 (excitatory and inhibitory; layers 2/3 and 5/6). We make connections to pyramidal cells in layer 5/6 comparatively stronger than those targeting pyramidal cells at layer 2/3, in line with anatomical data [2] and recent experimental data showing that top-down signals selectively activate infragranular layers in human [34]. This inter-areal circuit motif is displayed in Fig. 4A and D.

**Figure 4:**
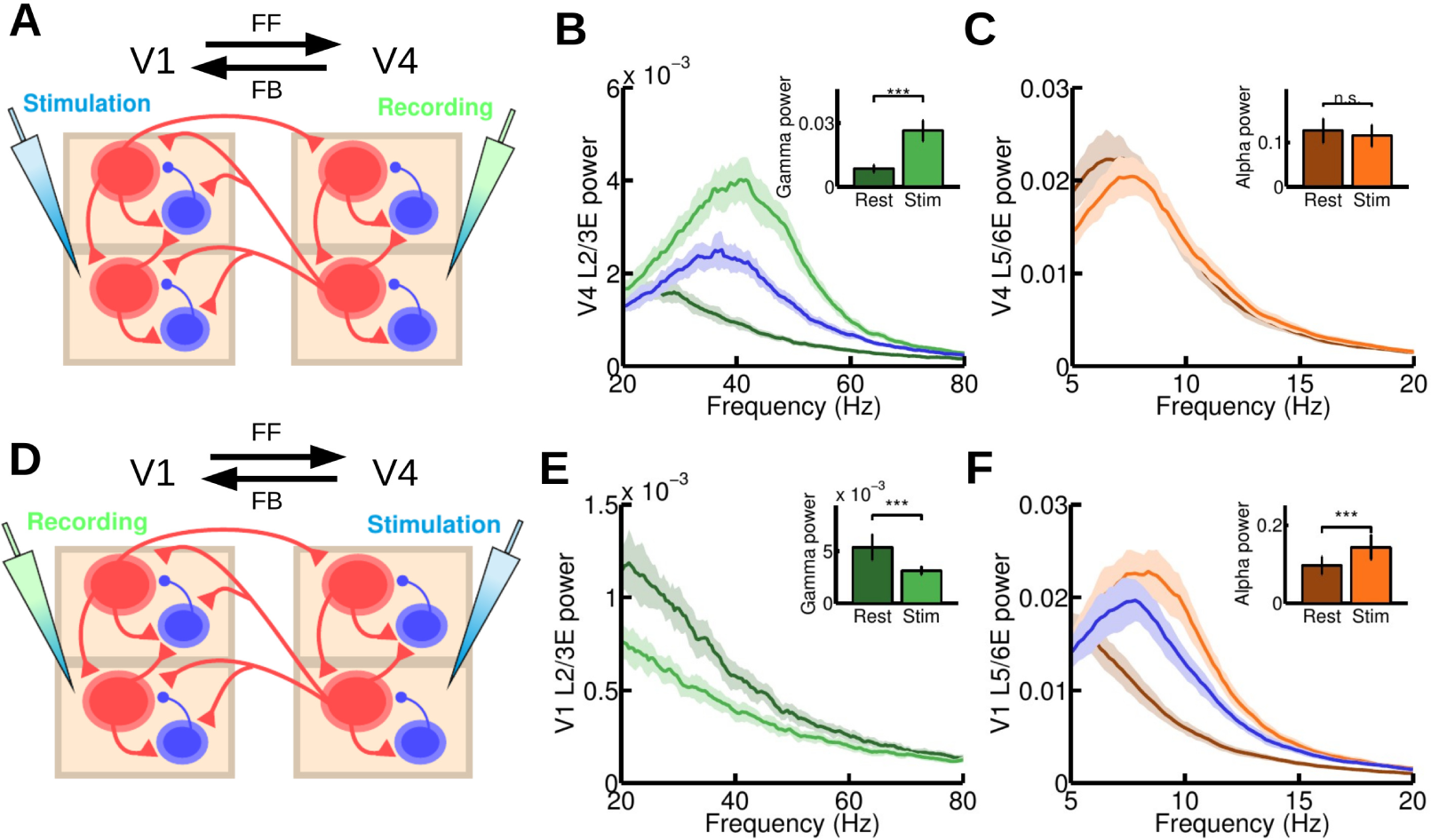
Microstimulation at the inter-areal level. (A) Scheme of the inter-areal projections between two areas (V1 and V4), the anatomical hierarchy ascends from left to right. We inject a current of I=15 at both supragranular and infragranular excitatory populations of V1 and measure at V4. A background current to excitatory populations in supragranular (I=2) and infragranular (I=4) layers in both V1 and V4 is injected to guarantee a minimum level of activity. (B, C) Power spectrum at V4 measured at layer 2/3 (B) and layer 5/6 (C), for resting and stimulation conditions. Insets show the (raw) peak value of the power spectrum at supragranular (B) and infragranular (C) layers for the same resting and stimulation conditions. A highly significant increase (p<0.001) in gamma power is found, as in the microstimulation experiments by van Kerkoerle et *al.* (D) Same as (A), but injecting a current of I=15 at all excitatory populations of V4 and recording in V1. A background current of I=1 to all excitatory areas in V1 and V4 is injected, to guarantee a minimum of activity. (E, F) Power spectrum at V1 layer 2/3 (E) and layer 5/6 (F), for resting and stimulation conditions. Inset shows the peak value of the power spectrum for these conditions. A highly significant increase (p<0.001) in alpha power and decrease (p<0.001) in gamma power were found, in agreement with experimental observations. For panels (B) and (F), the blue curve corresponds to an isolated area receiving the same input as in the stimulation case, but without its rhythmic component (see main text for details).

Inter-areal interactions in the visual areas have been recently studied in the context of visual attention [5, 4, 6]. Of particular interest for informing circuit mechanisms, van Kerkoerle et al. used microstimulation to reveal an enhancement of gamma/alpha power in FF/FB interactions [4]. With a model of two multi-laminar areas we can study these interactions and explore the underlying mechanisms involved.

We first simulated a purely FF communication (Fig. 4A): an injection of current into excitatory cells in V1 produced a highly significant increase in gamma power in layer 2/3 of V4 (Fig. 4B) and a small, non-significant decrease in alpha power in layer 5/6 of V4 (Fig. 4C). These effects closely resemble the results in [4] (cf. its figure 8a) and support the notion that strong gamma rhythms reflect FF communication. According to our model, this large increase is explained by two factors: the extra input arriving at V4 layer 2/3 leads to a mean-driven increase in gamma power similar to that observed in the local circuit (cf. Fig. 2B), and the gamma modulation of the arriving input further enhances the intrinsic V4 gamma rhythm via inter-areal synchronization. By considering V4 as an isolated multi-laminar area and injecting into its layer 2/3 a constant input equivalent to the one received from V1 in the two-area case, we can compare both effects (Fig. 4B). This reveals that a constant input enhancing the noise-driven gamma rhythm alone accounts for a major part of the observed increase in gamma power, although the effect exclusively due to inter-areal gamma synchronization (i.e. the difference between the blue and bright green curves in Fig 4b) makes a substantial contribution.

To simulate a purely FB communication, we injected current into excitatory cells in V4 and measured responses in V1 (Fig. 4D). This led to a drastic decrease in V1 layer 2/3 gamma power and a strong and significant increase in V1 layer 5/6 alpha power (Figs. 4E and F). Such behavior can be observed in the experimental microstimulation experiment in [4] (cf. its figure 8e), and suggests that strong alpha oscillations are associated with FB interactions (along with gamma rhythm suppression). This mechanistic explanation combines several factors: the alpha-modulated input arriving from V4 increases the alpha power in V1 layer 5/6 via (i) a larger average input (as in Fig. 2C) and (ii) the synchronization of both areas in the alpha range. As for FF, we can isolate area V1 and inject the equivalent constant input. This shows that the increase of alpha power due to factor (i) is much larger than the increase due to (ii). The decrease of gamma power is also observed experimentally in [4] and is in agreement with our findings at the inter-laminar level (Fig. 3C). This decrease is explained by the net inhibitory effect that V1 layer 5/6 exerts on V1 layer 2/3 plus the FB from V4 to layer 2/3 inhibitory cells, which overcomes the small excitatory contribution from V4 FB to V1 layer 2/3. Other possible scenarios, such as a strong FB connection projecting to supragranular layers, can be also considered in the model (see Fig. S1) and they provide interesting insights in the context of top-down attention signals (see Discussion).

The last scenario that we consider here is bidirectional communication, i.e. stimulating both areas V1 and V4 and analyzing the frequency-specific profile of the signals in each direction (Fig. 5A). The recorded signal for a given area is defined as a weighted combination of activity in layers 2/3 and 5/6, mimicking depth-electrode recording (see Supplementary Methods for details). By computing the spectral coherence between activity at V1 and V4, we observe two clear peaks located at the alpha/low beta and gamma ranges (Fig. 5B), in agreement with experimental recordings [4, 6] (cf. their figures 7b and 3d, respectively). We can further obtain information on directionality of signal flow by computing a frequency-dependent Granger causality (GC) analysis on both FF and FB directions. The results (Fig. 5C) show excellent agreement with experimental observations ([4, 6], cf. their figures 7d and 3b, respectively), supporting the notion that enhanced gamma and alpha/low beta rhythms are associated, respectively, with FF and FB communication. This result can also be easily quantified by defining, as in [6], the directed asymmetry index (DAI) between two areas as the (normalized) difference between spectral GC fluxes, i.e.

**Figure 5:**
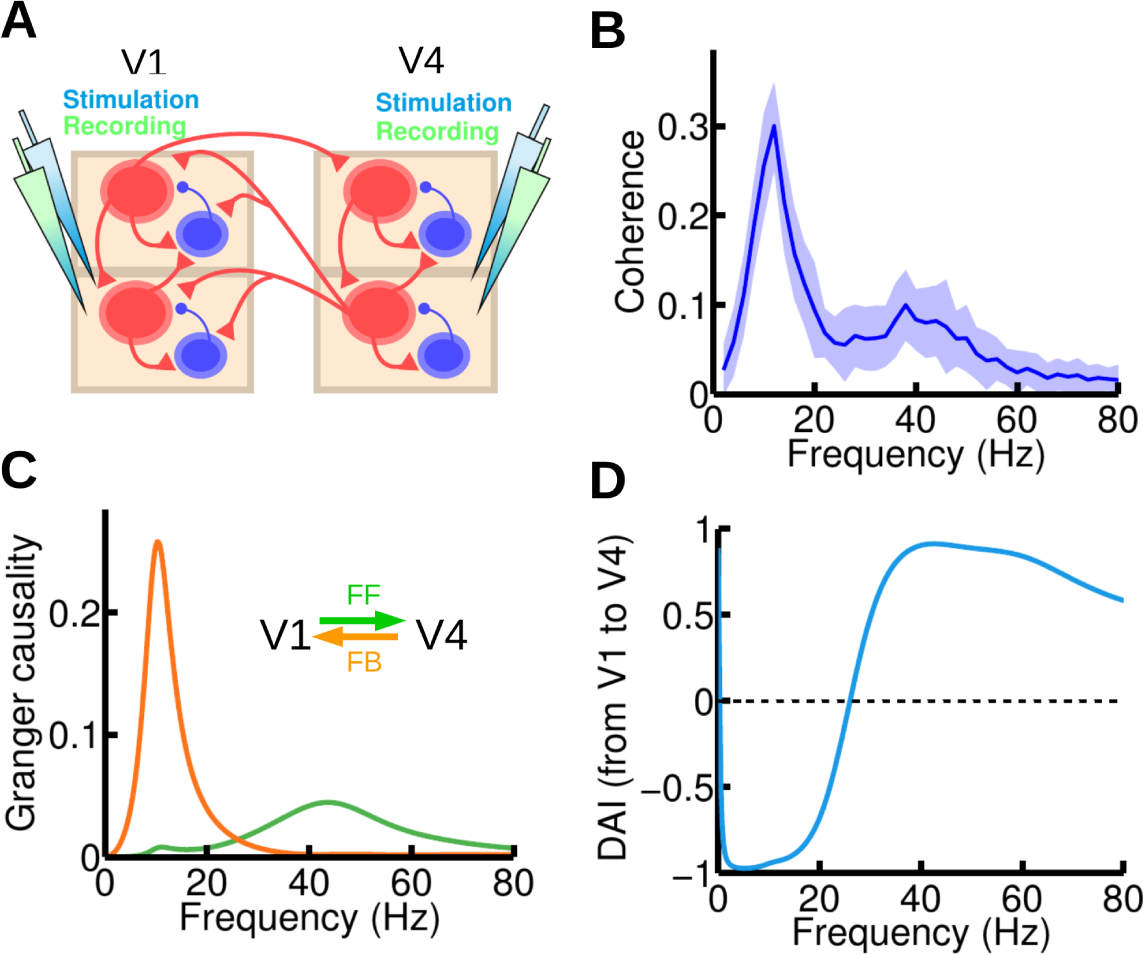
Frequency-specific feedforward and feedback interactions. (A) We now inject current and record the activity of both areas (V1 and V4). An input current of I=8 was injected in all excitatory populations of the circuit. (B) Spectral coherence between V1 and V4 activity highlights the existence of two peaks, at the alpha and gamma range respectively. (C) Spectral pairwise Granger causality in the V1-to-V4 (green) and V4-to-V1 (orange) directions, showing that each of the peaks found in (B) corresponds to a particular direction of influence, suggesting that frequency-dependent Granger-causality analysis could be used to deduce feedforward vs feedback signaling directionality between areas for which the hierarchical positions are not known anatomically. (D) The DAI profile of the functional connection, which is obtained by normalizing the difference between the two GC profiles in (C), can be used to characterize a directed functional connection between two cortical areas.

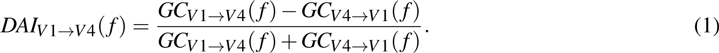

Note that 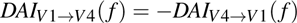. The DAI profile, which defines a frequency-specific directed functional connection, is shown for the example areas V1 and V4 in Fig. 5D.

In summary, our model of two interconnected cortical areas reveals frequency-dependent signatures in each direction as observed experimentally. Due to a paucity of available data, we adjusted in the model the relative weights of connections from a feedback projection onto the supragranular and infragranular layers, and found that the projection should be predominantly targeting the infragranular population in order for the model to reproduce the recent electrophysiological observations on the frequency-dependent feedforward versus feedback signaling. Combined with the connection from infragranular pyramidal cells to supragranular interneurons, this result suggests a net inhibitory influence of top-down signal which can potentially be compared with a bottom-up signal via feedforward excitation in the supragranular circuit.

### 2.4 Large-scale level

In order to develop our inter-areal system into a large-scale model, we make use of anatomical connectivity data from an ongoing tract tracing study in macaque [9, 11, 2]. In brief, retrograde tracer injected into a given target area labels neurons in multiple source areas that directly project to the target area. The proportion of labeled neurons in a given source area defines a weight index as the fraction of labeled neurons (FLN) from that source to the target area. Additionally, the number of labeled neurons located in the supragranular layer of a given source area (over the total number of labeled neurons in that area) defines the lamination index SLN (for fraction of supragranular layered neurons for a given pathway from source to target area). Source areas that are lower (higher) than the target area in the anatomical hierarchy tend to have a progressively higher (lower) SLN. In other words, the lower (higher) the source area relative to the target area, the higher (lower) the SLN values of the source-to-target projection (see Fig. 6A). For instance, purely FF or FB projections have SLN value of 1 and 0, respectively. By repeating the process using other anatomical areas as target areas, an anatomical connectivity dataset with weighted directed connections and laminar specificity is obtained (Figs. S2 and S3). Elsewhere we have shown that SLN captures the hierarchical distance so that individual SLN values allocate individual areas in the appropriate ranking according to the Felleman and Van Essen 1991 model [2, 1].

**Figure 6:**
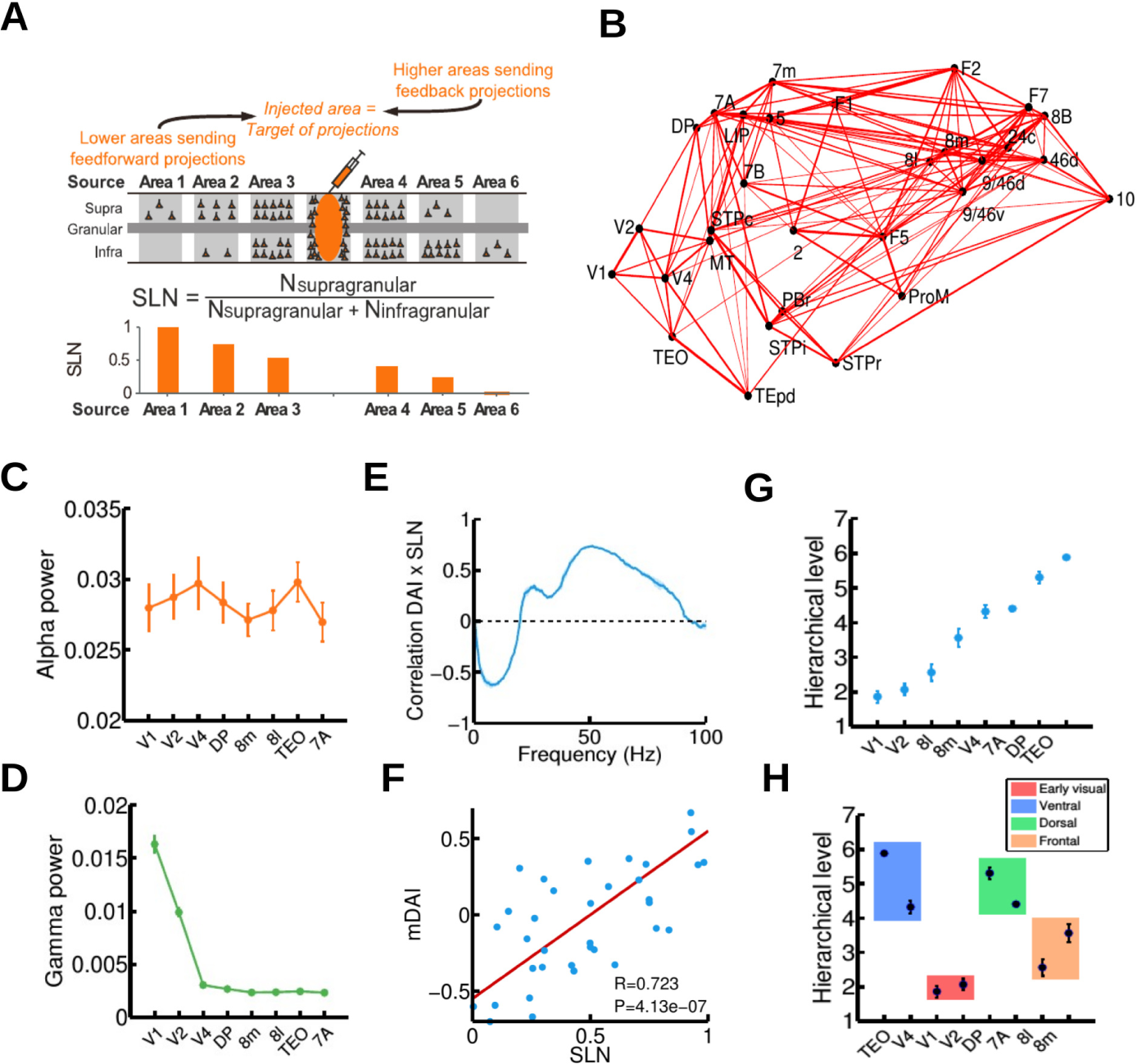
Large-scale cortical network and functional hierarchy. (A) Illustration of the anatomical tract tracing technique used to obtain the anatomical large-scale network, and in particular the fraction of supragranular labeled neurons (or SLNs, see main text for details). A high (low) value of SLN for a given projection indicates that the source area is lower (higher) than the target area (the injected area) in the anatomical hierarchy. (B) 3D plot of the macaque anatomical network obtained (only projections with FLN>0.005 are plotted, for clarity), with all 30 areas in their spatial positions. Connection strength is indicated by line width. (C, D) Alpha (C) and gamma (D) power for eight selected cortical areas of interest (V1, V2, V4, DP, 8m, 8l, TEO, and 7A). (E) Correlation between SLN and DAI, as a function of frequency. The correlation is positive in the gamma range and negative in the alpha range, indicating a prevalent involvement of these rhythms in FF and FB interactions, respectively. (F) Correlation between SLN and the combined DAI across gamma (30-70 Hz) and alpha/low beta (6-18 Hz) frequency ranges (named mDAI, see text for details). (G) Functional hierarchy emerging from the frequency-specific interactions in the network, and computed using the mDAI values as in Bastos *et al*. (H) Areas belonging to the same type (early visual, ventral, dorsal or frontal; indicated by color box) tend to be clustered in the same way as in the experimental observations. For all panels, visual input was simulated with an input current I=8 to the supragranular excitatory population of V1, and a background current of I=6 to all excitatory populations in the network was considered.

In the present study we use the recently acquired anatomical connectivity dataset [2], and in addition have performed additional injections to expand the connectivity data and include the parietal area LIP in the connectivity dataset. The new connectivity, displayed as a 3D network in Fig. 6B, has 30 cortical areas with a graph density of around 66% [3]. To extend our inter-areal to a large-scale model, we set a multi-laminar circuit at each node of this anatomical network and use (i) FLN values as an estimate of the weight or strength of the inter-areal connections, (ii) the estimated wiring distances between injection sites to model axonal propagation delays between cortical areas (Fig. S4), and (iii) the SLN values as a measure of hierarchical distance between areas.

To simulate an increased intensity of the visual stimulus, we increase the level of constant current arriving at V1 layer 2/3 neurons, and investigate the resulting interactions among cortical areas and their rhythmic activity. Figs. 6C and D show the alpha and gamma peak power observed in the model, for a subset of cortical areas (V1, V2, V4, TEO, 8m, 8l, DP, and 7A; their anatomical relations are shown in Fig. S5) whose interactions have been analyzed in the context of visual attention [6]. We observe a relative increase in the alpha rhythms in areas of the ventral stream (V4 and TEO). We also find an especially strong gamma rhythm in early visual areas (V1 and V2) due to sensory input, in agreement with observations of sensory-driven gamma rhythm enhancements being prominent in early visual cortex [7]. To characterize the frequency-specific interactions between areas, we compute spectral pair-wise conditional Granger causality (GC) profiles between the subset of areas, as in [6]. For all area pairs we observe a relationship between FF/FB interactions and gamma/alpha rhythms (Fig. S6) similar to that found earlier with our two-area model (Fig. 5C).

From the spectral GC for all areas, we obtain the spectral DAI profile for each area pair (as in our two-area model), which defines a directed spectral functional connectivity. By computing, for each frequency, the correlation between these functional connections and the SLN data (which provides information about the directionality of the anatomical projections), we find a clear pattern: SLN and DAI are positively correlated in the gamma range, and negatively correlated in the alpha/low beta range (Fig. 6E). This agrees with recent experimental findings [6], and it serves as a quantitative demonstration that the strong gamma/alpha signature of FF/FB interactions holds at the level of a dense, large-scale network. To further quantify this correlation, we define the multi-frequency DAI (or mDAI) per area pair as the averaged DAI for both gamma and alpha ranges, i.e.

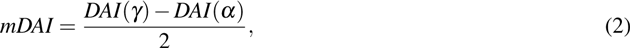

where the negative sign in the second term accounts for the negative correlation of SLN and DAI in the alpha range (see Supplementary Methods). Using this quantity, we obtain a highly significant correlation coefficient (Fig. 6F) between anatomical projections and functional interactions, in accordance with experimental studies [6].

The laminar pattern of anatomical FF and FB projections is the defining feature of the global anatomical hierarchy of visual areas [1, 2]. Since our model displays a strong correlation between anatomical projections and functional interactions, it is interesting to test whether the model predicts the emergence of a similar hierarchy at the functional level, as recently observed in vivo [6, 7]. We follow the same procedure as in [6] to define the hierarchical distance between area pairs from mDAI values (see Supplementary Methods), and after simulating the full large-scale model and computing its mDAI values, we observe the emergence of a clear functional hierarchy among visual areas, as shown in Fig. 6G. As in the experimental functional hierarchy [6], early visual areas lie at the bottom of the hierarchy, followed by areas of the frontal eye field (8l and 8m) and with extrastriate visual areas of the ventral and dorsal functional streams at the top. Areas within the same functional streams are clustered around similar hierarchical values (Fig. 6H), in agreement with [6]. These results show that the spectral functional interactions, as well as the formation of a functional hierarchy observed experimentally, can be explained within a computational framework of locally generated rhythms propagated through a multi-laminar network structure.

As an interesting example of the predictive power of our large-scale model, we use it to analyze a complex phenomenon observed during visual attention tasks. It has been reported that, contrary to the anatomical hierarchy, the functional hierarchy is not fixed. More precisely, Bastos et al. found that the position of visual areas in the functional hierarchy are highly dynamic and switch locations in a context-dependent fashion [6]. The ranking of areas in the functionally defined hierarchy in the pre-cue period of the task differ significantly from the ranking obtained in the post-cue period, when top-down modulations from higher areas are expected to heavily influence visual areas at the bottom of the hierarchy. These “hierarchical jumps” were especially noticeable in frontal eye field areas, such as 8l and 8m, and their origin and implications are as yet unknown.

To provide a mechanistic explanation for jumps in functional hierarchy, we first analyze a simple network of a multi-laminar area pair with a well-defined anatomical hierarchical relationship (Fig. 7A). In this scenario, the functional hierarchical distance between both areas is given by the mDAI value of the pair, which can be decomposed into the DAI in the gamma and alpha ranges (see Eq. 2). Considering the spectral DAI in the ascending anatomical direction, the more positive the DAI in the gamma range (and the more negative in the alpha range), the larger the hierarchical distance. Since external input to specific layers modifies the gamma and/or alpha power in a given area, it is plausible that a laminar-specific input could affect the frequency-specific interactions between areas and therefore their hierarchical distance. To test this hypothesis we measured the frequency-specific interactions between both areas for three types of laminar input patterns to the lower area: (i) a small input to layer 2/3 and a large input to layer 5/6, (ii) an equal input to both layer 2/3 and 5/6, and (iii) a large input to layer 2/3 and a small input to layer 5/6. Changes in the laminar-specific inputs have a substantial impact on the DAI profile, increasing the DAI in the gamma range (and therefore the FF interactions) with increases in the supragranular-to-infragranular input ratio (Fig. 7B). Such gamma enhancements are due to the combined effect of the increase in direct excitation to layer 2/3 and the decrease of inter-laminar inhibition from layer 5/6. Increases of DAI in the gamma range imply a higher mDAI value of the area pair (Fig. 7C), leading to an increase in the hierarchical distance for this simple network.

**Figure 7:**
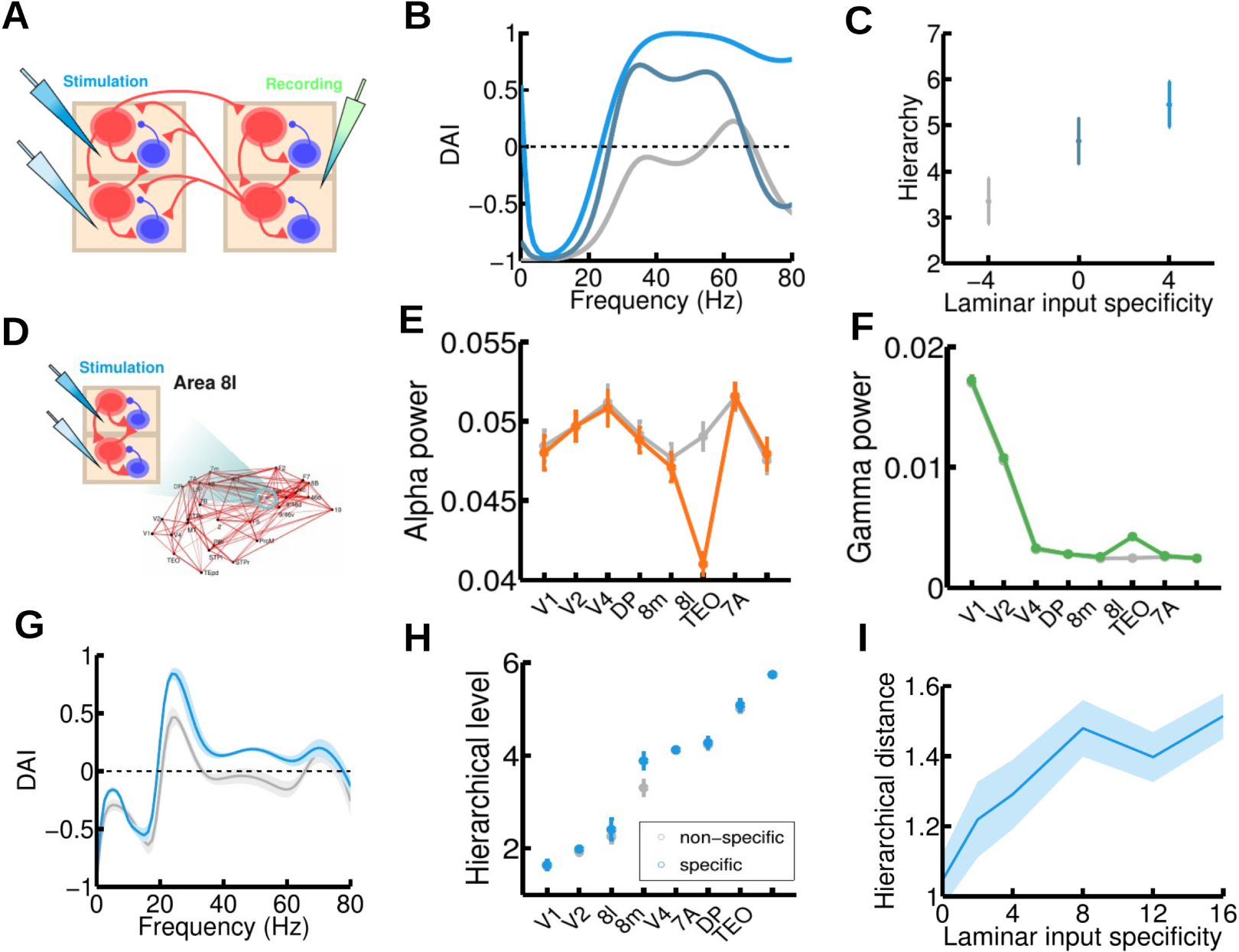
Mechanistic explanation for the experimentally observed hierarchical jumps. (A) Scheme of the simple two-area circuit considered (we use the same parameters as for the two-area microstimulation protocol); the area on the left, which is lower in the anatomical hierarchy, receives laminar-specific input. (B) The gamma component of the DAI increases as the input to layer 2/3 exceeds input to layer 5/6 in the lower area (ascending curves correspond to input to L2/3E increasing from I=4 to 6 to 8, and input to L5/6E decreasing from I=8 to 6 to 4). Excitatory populations in the higher area receive a fixed I=6 background current. (C) Increases in the hierarchy rank of the higher area as a consequence of the laminar-specific input in (B). The laminar specificity, S, is defined as the difference between the input to L2/3E and to L5/6E. (D) We follow the same procedure, but now injecting laminar-specific current into area 8l within the full 30-area network. (E, F) Changes in alpha (E) and gamma (F) band power as a consequence of the injection of laminar-specific input. Lines in gray correspond to the same input injected at both layers (i.e. no laminar-specific input), while colored lines correspond to a laminar specificity of S=8. (G) The spectral DAI profile from 8l to 8m increases in the gamma range as a consequence of the laminar-specific input (grey curve: S=0, blue curve: S=8). (H) A hierarchical jump of area 8m is observed, as in the two-area case (gray points: S=0, blue points: S=8). (I) We find a robust increase of the hierarchical jump distance with the strength of the laminar specificity of the input. Other parameters and background currents are as in Fig. 6

Extending this analysis to the full 30-area network is not a straightforward process, since cortical areas tend to interact in a non-trivial fashion and changing the laminar specificity of the input to one area influences interactions with several areas simultaneously. We can, however, test our hypothesis in a concrete case that is of particular interest. According to the FLN data (see Fig. S2), areas 8l and 8m are strongly connected to each other, while their projections to and from other areas are weaker. This provides a useful testbed for translating our findings from the case of two isolated areas to the full cortical network. In terms of laminar origin, projections between areas 8l and 8m are approximately horizontal (i.e., SLN is about 0.5 in both directions, see Fig. S5). Furthermore, both areas display a high hierarchical mobility according to experimental observations [6], so the pair 8l-8m is a suitable candidate for observing hierarchical jumps in our model.

Since 8l is lower than 8m in the functional hierarchy (both in the model and in the experimental findings), we simulate the large-scale network of 30 areas with a highly laminar-specific input to area 8l (strong for layer 2/3, weak for layer 5/6, see Fig. 7D), and compared the DAI profiles and functional hierarchy with the original simulation (for which the input to 8l does not have laminar specificity). This laminar specific input elevates the gamma power and decreases alpha power in area 8l, as expected (Figs. 7E and F). Furthermore, Fig. 7G shows an important effect in the spectral DAI between 8l and 8m in this situation: the DAI from 8l to 8m is stronger in the gamma range, as in the two-area case (although, due to the interactions of other areas, the profile is more complex in this case). The main effect in the functional hierarchy (Fig. 7H) is an elevation of area 8m, which effectively increases the hierarchical distance between 8m and 8l as expected and induces a hierarchical jump of area 8m, as experimentally observed in [6]. To further test this effect, we repeat the process for several degrees of laminar specificity of the 8l input, and we observe in Fig. 7I a clear increase in the hierarchical distance between 8m and 8l as the laminar specificity of the input increases (stronger for layer 2/3 and weaker for layer 5/6).

These results indicate a strong prediction of our large-scale model: jumps observed in the dynamic functional hierarchy may reflect a change in the laminar specificity of the input to cortical areas. Such a change in the laminar pattern of the input could be due, for instance, to context-dependent influences from higher cortical areas, known to be important in attention and other cognitive tasks.

## 3 Discussion

The brain is characterized by interconnectivity and dynamics across many scales. Perhaps no recent experimental finding better highlights the challenge raised by this multi-scale complexity than the layer-specific, and frequency-dependent, interplay between feedforward and feedback signaling streams across the large-scale primate cortical circuit [5, 4, 6, 7]. To uncover the circuit mechanism behind these frequency-dependent processes, and to understand their implications for large-scale communication, we have built a multi-scale model of the macaque brain covering both slow (alpha) and fast (gamma) neural oscillatory dynamics and four spatial levels of description constrained by the known anatomy. The spatial levels range from local homogeneous populations to laminar circuits, inter-areal interactions, and large-scale cortical networks based on precise anatomical macaque connectivity data. The parsimonious model is able to explain a wide range of macaque electrophysiological observations, including the effects of visual contrast on V1 gamma rhythms [13, 14], the inter-laminar phase-amplitude coupling between gamma and alpha rhythms [5, 16], the relationship between alpha power and local inhibition [18], gamma/alpha signatures of FF/FB inter-areal interactions [4, 6], and correlations between anatomical and functional networks [6]. Notably, using the same analysis as in the experimental studies, our model reveals the emergence of a dynamic functional hierarchy in the visual cortex [6, 7].

The model makes a number of experimentally testable predictions. At the inter-laminar level, we suggest that the phase-amplitude coupling between gamma and alpha rhythms is mediated by projections from layer 5/6 pyramidal neurons to layer 2/3 interneurons. This sets a clear infragranular-to-supragranular direction of the modulation (as tentatively discussed in [16]) and highlights a key role of layer 2/3 interneurons in phase-amplitude coupling. Physiological experiments have in fact identified an inhibitory subgroup (including chandelier cells and irregular-spiking basket cells) as the recipient of strong projections from layer 5 pyramidal neurons [30, 31]. Our model suggests that this subgroup of interneurons will play an important role on phase-amplitude coupling between supragranular gamma and infragranular alpha rhythms, a prediction that could be experimentally tested.

The relevance of layer 2/3 interneurons on phase-amplitude coupling is expected to have important implications at the functional level. Since enhanced gamma rhythms are associated with a strong drive to higher order areas [4, 6], modulating the phase-amplitude coupling could lead to a modulation of the temporal windows of elevated gamma synchrony. Therefore, top-down signals targeting layer 2/3 interneurons (which are compatible with the unspecific FB projections considered in our model) could act as a top-down gating mechanism, controlling the length and occurrence of the temporal windows for communication [35].

At the inter-areal level, our model provides insight into the mechanisms that subserve cortico-cortical communication [36]. For FF communication, we have identified two mechanisms that may contribute to the experimentally observed enhancement of gamma power [4, 6]: (i) mean-driven enhancement of oscillations, and (ii) inter-areal synchronization. Both mechanisms contribute to the observed gamma rhythm enhancement, but due to the existence of the mean-driven enhancement mechanism, a tight synchronization between areas is not necessary to explain the experimental observations. Gamma rhythms tend to be very weakly coherent and a communication mechanism purely based on inter-areal gamma synchronization could pose certain problems [14, 37] (but see [36]). However, the presence of a certain level of inter-areal synchronization clearly enhances the gamma power associated to FF interactions, and its effect is stronger than the contributions of synchronization to FB interactions.

Although point-to-point FB projections between supragranular layers [2] are not explicitly incorporated into the model, inter-areal projections between supragranular layers in the FB direction are present in the model once we incorporate the anatomical data into the framework. For instance, a projection with an SLN value of 0.2 will be mainly a FB projection, but since the SLN value is larger than zero, the projection will have a (comparatively weak) supragranular-to-supragranular component to reflect the small fraction of supragranular neurons anatomically identified for that specific projection. Since weak but nonzero values of SLN are common in our anatomical dataset, this tends to be the norm rather than the exception, and the effect is the appearance of small peaks of GC at gamma ranges for projections that are generally considered as FB (Fig. S6). A more careful consideration of point-to-point FB projections from supragranular layers, however, is beyond the scope of this work, as it would involve extending the model of cortical area to include spatial extension and lateral connectivity within nearby neurons. Such extension will be considered in future studies.

The anatomical connectivity data used at the large-scale level contains detailed information about 30 cortical areas of the macaque brain [9, 2]. Other brain areas, including thalamic and other subcortical structures, are currently missing from the dataset and have therefore not been explicitly considered in the present model. While it is not uncommon to assume, as we do in this study, a stereotypical thalamic input to canonical cortical microcircuits [9], the absence of an explicit thalamic model may be seen here as an opportunity to conceptually explore the role of thalamus in cortico-cortical communication. For instance, a strong sustained input to all areas is needed in our model to guarantees that both gamma and alpha rhythms are sustained in the cortex, and such an input would be most likely provided by the thalamus. In addition, thalamic input could be also partly responsible of modulating the interactions between cortical populations, and therefore influencing the phase-amplitude coupling between cortical layers, for instance.

One could assume that dynamic thalamo-cortical interactions, or other mechanisms such as bursting, are necessary to generate cortical rhythms, as it is commonly assumed for alpha oscillations [38, 39]. While considering more detailed mechanisms of alpha rhythms is an appealing future direction in this context (and several modeling works could constitute a solid starting point, see [40, 38, 39]), it is unlikely that such mechanisms would significantly alter the conclusions of the present work. This is because, while further detail can be added into the rhythm generator, model predictions on power and frequency modulations induced by external input are consistent with the experimental observations. As a consequence, the transmission of signals between cortical areas and the modulation of these signals will accurately reflects the electrophysiological evidence available, independently of the particular mechanism for rhythm generation assumed for the model.

### 3.1 Predictive coding and selective attention

Synaptic connections at different levels of the model have been established according to known anatomical patterns [20, 24, 9, 2]. Full mapping of the model connections with anatomy is not presently possible, in part due to simplifications needed for a computationally tractable large-scale cortical model and partly to the current limitations in the anatomical literature. In particular, while the laminar origin of inter-areal projections is quantitatively measured for each pair of areas in our network [2], the laminar target of these projections can only be grossly estimated from other anatomical studies [41, 1, 2]. We have assumed, for example, that FF connections arriving to a given area exclusively target its supragranular laminar module, based on the anatomical evidence that layer 4 is the major target of FF projections [41, 1, 2]. There is anatomical evidence that in inter-areal pathways, supragranular layers target supragranular layers and infragranular layers target infragranular layers [42, 2], which supports this assumption. Interestingly, there is also evidence suggesting that FF projections could also aim at infragranular layers as secondary targets [43, 44]. This novel FF pathway would not substantially affect our modeling results, since the addition of such a secondary target for FF projections could be incorporated, as a first approximation, as a small increase in the strengths of the FF projection to layer 2/3 and the inter-laminar projection from layer 2/3 to layer 5/6.

The case of the laminar target of FB projections is arguably more interesting. It is thought that FB projections preferentially target supragranular and infragranular layers while avoiding layer 4 [2, 42]. The precise weight of the FB input to infragranular layers in early areas, however, is not known. In our model, we set connections to pyramidal cells in layer 5/6 comparatively stronger than those targeting pyramidal cells at layer 2/3, motivated by anatomical studies [2, 42]. As a result, FB projections in our model are net excitatory for layer 5/6 and net inhibitory for layer 2/3 (due to FB projections to interneurons in layer 2/3 plus the FB-activated local ascending inhibition from layer 5/6). Interestingly, such a connectivity pattern would be consistent with a predictive coding framework, since top-down signals would provide the strong inhibitory input to layer 2/3 necessary to suppress any predicted signals [45, 19]. In such a framework, only ascending information that could not be predicted and matched by the top-down input would form prediction errors able to move forward in the hierarchy through subsequent supragranular layers. This important role of strong inhibitory top-down signals on predictive coding, which has been previously discussed [19, 46], fits in our large-scale model in a natural manner.

Despite the general net inhibitory effect of FB projections, the existence of top-down excitatory interactions [41] may potentially play an important role in other contexts, such as in selective attention [47, 48]. As discussed in [19], predictive coding and selective attention are not mutually exclusive, and the potentiation of behaviorally relevant bottom-up signals through top-down attentional modulation is also an important function in a predictive coding scenario. FB projections able to provide a net excitatory contribution to supragranular layers have been anatomically identified [41, 2], and our model is able to shed light as to its potential role in selective attention. In particular, by considering the existence of net excitatory FB projections to layer 2/3 (and leaving the FB projections to pyramidal neurons of layer 5/6 as a comparatively weaker projection), our model predicts a decrease of infragranular alpha power and an enhancement of gamma rhythms and supragranular firing rate induced by top-down signals (see Fig. S1). In this case, top-down input to layer 2/3 significantly enhances gamma power by increasing the level of excitation in the circuit, with inter-areal synchronization playing only a minor role. These results are in agreement with attention-related V4 activity recorded from macaque, where similar gamma/alpha signatures are observed in receptive fields covering the attended visual stimulus [47, 49]. We suggest that a routing mechanism at higher cortical areas should be able to selectively activate either of the two FB pathways (the net inhibitory one or the net excitatory one) to prevent or facilitate the propagation of bottom-up signals in a context-dependent manner.

### 3.2 Relation to other models

It is useful to discuss how our model compares to other laminar models of a local cortical area. Kopell and collaborators, for instance, developed a laminar spiking model with three layers and different types of interneurons, which provides interesting insight on the top-down attention mechanisms mentioned above [50], and also on the role of cholinergic neuromodulators in inter-areal communication [51]. Diesmann and collaborators have also developed an anatomically-and physiologically-based laminar spiking model of a cortical microcolumn [33], with connectivity patterns in agreement with our simplified laminar circuit. Our approach of considering a parsimonious model with two laminar modules has several distinctive features and advantages at the conceptual level (for example, it provides a cleaner picture to understand the origin of phase-amplitude coupling) and in the context of scaling up the model to large-scale networks. Additional features from the above or similar laminar models can be incorporated in our large-scale framework to further increase the multi-scale predictive power of our approach in the future.

At the large-scale level, recent models of the macaque brain have highlighted the importance of heterogeneity to explain the emergence of a hierarchy of time scales [52], as recently observed experimentally [53, 54], although these models lack structure at the laminar level. Including a certain level of heterogeneity in a laminar large-scale model could reveal new mechanisms for inter-areal communication. It will be worth examining, in future research, the incorporation into our new model of different types of heterogeneity (in particular, leading to different gamma frequencies [14, 36]) that may contribute to efficient coordination of global brain activity. This is particularly interesting since, although cell-to-cell heterogeneity is known to have a strong effect in the dynamics of local populations [55, 56], the effect of area-to-area heterogeneity on large-scale networks is much less understood.

### 3.3 Functional hierarchy

Whereas the brain hierarchy is robust, it is also flexible functionally. A strong and layer-specific input is able to modify the functional hierarchical distance between cortical areas and to induce “jumps” in the hierarchy (see Results). This jumping phenomenon may result from layer-specific input arrives from cortical areas higher in the hierarchy, such as parietal or prefrontal areas. In particular, areas involved in higher cognitive functions such as working memory or rule representation should be able to send layer-specific top-down signals in a context-specific manner, leading to a reorganization of sensory and association areas in the functional hierarchy as observed experimentally [6]. The thalamocortical system may also contribute to altering brains functional hierarchy, a possibility that can be analyzed in our large-scale model extended to include the thalamus.

Finally, understanding the cortical mechanisms underlying hierarchical processing will improve future estimates of the anatomical hierarchy in humans. The measurement of laminar-specific projections via tract tracing studies defines an anatomical hierarchy in macaque [1, 2], but this anatomical data is not available with known imaging or postmortem technique in humans. Diffusion tensor imaging does not provide directionality information about inter-areal connectivity. Therefore, anatomically defined hierarchy is not known for humans at the present time. However, inspired by studies showing a strong correlation between functional hierarchy obtained from frequency-dependent Granger-causality analysis with anatomically defined hierarchy, a recent human study showed that the same analysis applied to magnetoencephalography yielded a functional hierarchy in humans [7], allowing inference of anatomical hierarchy of the human cortex at least for areas showing strong homology with macaque. Computational modeling studies such as the one presented here will contribute importantly to solve this problem and will strengthen the links between functional and anatomical connectomes.

## 4 Materials and Methods

### 4.1 Anatomical data

The anatomical connectivity data used in this work comes from an ongoing track tracing study in macaque [9, 11, 2]. In short, retrograde tracer is injected into a given target area and it labels neurons in several source areas projecting to the target area. The number of labeled neurons on a given source area allows to define the fraction of labeled neurons (FLN) from that source to the target area. Additionally, the number of labeled neurons located on the supragranular layer of a given source area (over the total number of labeled neurons on that source area) allows to define the supragranular layered neurons (SLN) from that source area to the target area. Source areas that are lower (higher) than the target area in the anatomical hierarchy tend to have a progressively higher (lower) proportion of labeled neurons in the supragranular layer. In other words, the lower (higher) the source area relative to the target area, the higher (lower) the SLN values of the source-to-target projection. By repeating the process using other anatomical areas as target areas, an anatomical connectivity matrix with weighted directed connections and laminar specificity can be obtained.

We use the anatomical connectivity matrix from recent studies [2], and we also perform additional injections to expand the connectivity data and include the parietal area LIP in the connectivity matrix. The 30 cortical areas, which constitute the new connectivity matrix, are: V1, V2, V4, TEO, 9/46d, F5, 8m, 7A, DP, 2, 5, 7B, STPr, STPi, STPc, PBr, TEpd, 24c, F1, F2, F7, ProM, 8L, 9/46v, 46d, 8B, MT, 7m, 10 and LIP.

The corresponding 30x30 matrices of FLN, SLN and wiring distance are shown in Fig S2, S3 and S4, respectively. These matrices are also shown in Fig S5 for a particular subset of cortical areas of interest (used for the study of the functional hierarchy in the model). The subset of areas of interest is V1, V2, V4, DP, 8m, 8l, TEO and 7A.

Surgical and histology procedures were in accordance with European requirements 86/609/EEC and approved by the ethics committee of the region Rhone-Alpes.

### 4.2 Computational model

The large-scale model of the macaque cortex is built by assembling four different levels of description of increasing size: (i) local excitatory-inhibitory populations of the Wilson-Cowan type that describe the activity within a given layer, with quantitatively differences for the supragranular and infragraular layers so that they preferentially exhibit noisy oscillations in the gamma and alpha frequency-band, respectively, (ii) inter-laminar circuits coupling supragranular and infragranular layers, (iii) inter-areal couplings which consider the layer-specific influences between two given cortical areas (such as V1 and V4), and (iv) large-scale laminar cortical network, which extends the anatomical connectivity data from [9, 2] to include the posterior parietal area LIP and a laminar structure of local areas. The network model consists of 30 cortical areas distributed among occipital, temporal, parietal and frontal lobes. Further details of the model, including equations, parameter values and the routines used for data analysis can be found in the Supplementary Materials.

## 5 Acknowledgements

This work was supported by the ONR grant N00014-13-1-0297, NIH grant R01MH062349, STCSM grants 14JC1404900 and 15JC1400104 (XJW) and by ANR-11-BSV4-501 and LABEX CORTEX (ANR-11-LABX-0042) of Université de Lyon, program “Investissements d’Avenir” (ANR-11-IDEX-0007) operated by the French National Research Agency (HK). Author contributions: JFM, JDM and XJW conceived the study, JFM performed the simulations, JFM, JDM and XJW analyzed the results, HK contributed data, and all authors contributed to the writing of the manuscript. Competing interests: The authors declare that they have no competing interests. Data and materials availability: Anatomical data can be downloaded from core-nets.com. The code with the computational model will be made freely available upon publication of the manuscript.

